# Multi-scale annotations of chromatin states in 127 human cell-types

**DOI:** 10.1101/2020.12.22.424078

**Authors:** Yan Kai, Stephanos Tsoucas, Shengbao Suo, Guo-Cheng Yuan

## Abstract

Genome-wide profiling of chromatin states has been widely used to characterize the biological function of non-coding genomic sequences in a cell-type specific manner. However, the systematic, comprehensive annotations of chromatin states from experimental data are challenging and require not just extensive biological knowledge but also sophisticated computational modeling. Previously we developed a hierarchical hidden Markov model, named diHMM, to systematically annotate chromatin states at multiple scales based on the combination of histone mark and chromatin regulator binding profiles. Here, we have improved the method by optimizing computational efficiency and using an ensemble-clustering approach to achieve a unified annotation by integrating information from cell-type-specific models. We then applied this improved method to generate a unified multi-scale chromatin state map in 127 human cell types, based on public data generated by the Epigenome Roadmap and ENCODE consortia. We found cell types with similar origin are typically associated with similar chromatin states, but cultured cell lines have distinct structures than primary cells. The contribution of enhancer elements to gene regulation is mediated by the broader context of domain-state organization. Distinct domain-state patterns are associated with various 3D chromatin structures. As such, we have demonstrated the utility of the multi-scale chromatin state map in characterizing the biological function of the human genome.

## Background

In eukaryotic cells the genome sequence is wrapped around the chromatin, with the nucleosome as its fundamental unit. Each nucleosome is an octamer of histone proteins, which, in addition to packaging the chromatin, can also play an important role in regulating transcriptional activities by forming a combinatorial chromatin state which includes covalent modifications, histone variants, and forming higher order structure [1]. While the underlying genome sequence is almost identical from cell to cell, the chromatin states can be highly variable. Such variations are essential for maintaining the diversity of gene expression programs, therefore the functionalities, across different cell types within the same organism.

Extensive efforts have been dedicated to map the chromatin states in different cell types (for example, by the International Human Epigenome Consortium and affiliated members). Among these efforts, computational method development has played an important role [2–5]. This is because it is a daunting task to systematically characterize the functionality of a huge number of possible patterns established by the combination of many signals on a single nucleosome by experimental methods alone.

Many methods have been developed over the years [2–5]. However, almost all these methods are designed for chromatin state characterization at a fixed length scale. Recognizing its limitation to study the hierarchical organization of chromatin, we have recently developed a hierarchical hidden Markov model, called diHMM [6], to simultaneously annotate chromatin state organization at multiple length scales. Using this model, we have demonstrated that the function of a DNA element depends not only on its chromatin state at its locus but also on the broader spatial context. However, due to the limitation of its computational efficiency, we were not able to use this method to compare the differences between multi-scale chromatin states across many cell types.

Here, we have significantly improved the computational efficiency by optimizing computational efficiency and using an ensemble-clustering approach to achieve a unified annotation by integrating information from cell-type-specific models. This allowed us to analyze two large public datasets generated by the ENCODE and Epigenome Roadmap consortia, spanning across 127 human cell-types, thus providing a comprehensive multiscale chromatin state reference map. To demonstrate its utility, we analyzed the variation of chromatin states across cell types and compared gene expression and 3D chromatin data and gained new insights.

## Results

### An improved diHMM algorithm annotates multi-scale chromatin states in 127 human cell-types

Our original implementation of the diHMM method was limited by computational inefficiency. It took about one week’s computer time to construct a model from only one chromosome from three cell lines. Therefore, it was challenging to annotate a large number of cell types. We have made a number of improvements to overcome this limitation. First, we implemented the diHMM algorithm in C++ by using a number of computationally efficient algorithms. This has improved the computational efficiency by ~30 times (see **Methods** for details). However, it is still infeasible to simultaneously annotate a large number of cell types, which would require a large amount of computing resources (CPU and memory) that we do not have access to.

As an alternative to building a single unified model by using all the data as the training set, we divided the task into smaller pieces by generating a number of cell-type specific models, each trained from a single cell-type. These models are slightly different due to the variation of training data, therefore their output cannot be directly compared. To build a unified model, we used a consensus clustering approach [7] to combine information from these cell-type specific models (**Supp Figure 1A**), which includes three major steps: (1) A diHMM model is trained for each cell-type individually; (2) Each model is applied to annotate all the cell-types, resulting in multiple annotations for each cell-type; and (3) A consensus set of annotation is constructed to maximize the agreement with the individual annotations (**Supp Figure 1B**, see **Methods** for details). Since each individual model is trained based on information from a single cell-type, the overall computing time and memory usage is vastly decreased and the process can be parallelized for further improvement of efficiency. The results are referred to consensus diHMM to distinguish from those obtained from applying diHMM to train a model from the whole dataset.

We applied the consensus diHMM approach to annotate the chromatin states in 127 cell types profiled by the Epigenome Roadmap [8] and ENCODE consortia [9]. ChIP-seq experiments were used to profile 13 histone marks and chromatin associated regulators, including H3K4me1/2/3, H3K27ac, H3K9ac, H3K79me3, H4K20me1, H3K36me3, H3K9me3, H3K27me3, H2A.Z, DNase hypersensitivity, and CTCF. Due to incompleteness of the data, we performed analysis not on the raw data directly, but on the imputed and binarized generously provided to us by Dr. Jason Ernst (see Data sources). To test if the consensus diHMM approach indeed improves agreement, we evaluated the normalized mutual information between the annotations obtained from the consensus diHMM and cell-type specific models, and found that the overall agreement was indeed improved (**Supp Figure 2**).

We used a similar strategy as in our previous work [6] to functionally annotate the consensus diHMM states, using information from the combinatorial pattern of ChIP-seq signals (**Figure 1B**), the enrichment of each chromatin state label in functional elements (**Supp Figure 3A**), and the fraction of genome covered by each state in the concatenated genome of 127 cell-types (**Supp Figure 3B**), and the spatial distribution around Transcription Start Sites (TSSs) (**Supp Figure 3C**). As before, the 30 nucleosome-level states were grouped into 14 broad categories (left panel in **Figure 1B**). Our nucleosome-level chromatin state interpretation and annotation are highly consistent with previous annotation efforts such as chromHMM [2] (**Supp Figure 4**).

**Figure 1.**
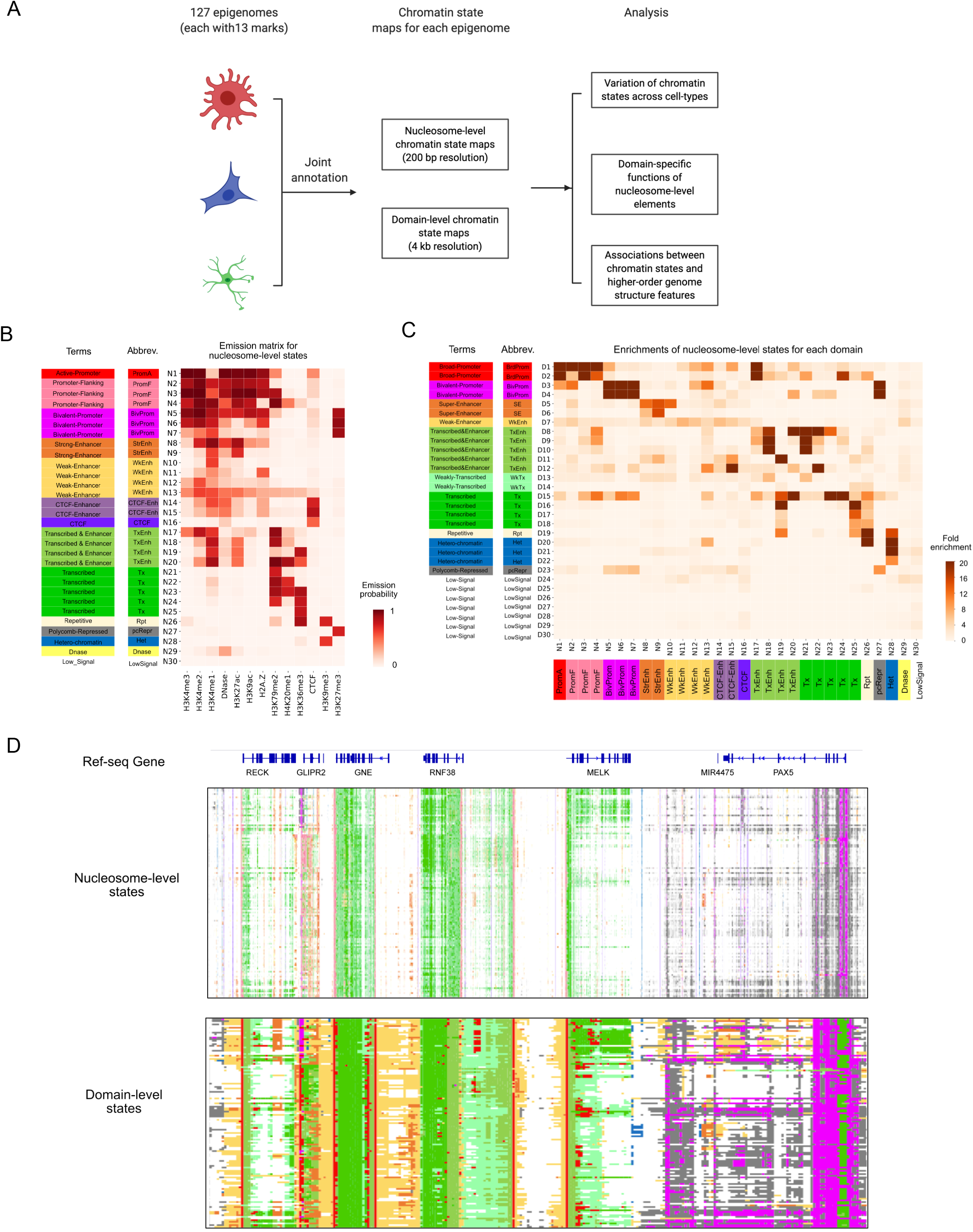
An overview of the multi-scale chromatin state annotations in 127 cell types. (A) A schematic overview of our analysis pipeline. (B) The combinatorial histone modification pattern associated with each nucleosome level chromatin state. Bars at the left show the names of functional terms and their abbreviations for the 30 nucleosome-level states. (C) Heatmap showing the enrichment of nucleosome-level states in the domain-level states. Bars at the left show the names of functional terms and their abbreviations for the 30 domain-level states. (D) A snapshot of the diHMM state annotations across different cell types. Each chromatin state is color-coded as in (B) and (C).

The domain-level chromatin states were annotated primarily based on their enrichment in the nucleosome-level states (**Figure 1C**), as well as the overall state coverage (**Supp Figure 3D**) and spatial distribution around TSSs (**Supp Figure 3E**). As a result, the 30 domain-level chromatin states were grouped into 11 broad categories. Taken together, we have generated a unified, multi-scale chromatin state map for the 127 human cell types. A snapshot of this map is illustrated in **Figure 1D**.

### Analysis of chromatin state variations across human cell types

We next sought to understand the overall similarity of the nucleosome- and domain-level chromatin state maps across the 127 chromatin state maps. We used the normalized mutual information (NMI) metric to quantify the similarity between different chromatin state maps in the same way as described in the previous section, and converted the results as a distance matrix (see **Methods**). For visualization, the t-distributed stochastic neighbor embedding (t-SNE) algorithm [10] was used to project the high-dimensional profile into a 2D space (**Figure 2A-B**). As expected, cells with similar lineage were projected next to each other, such as T-cells, B-cells, embryonic stem (ES) cells and heart cells. At a higher resolution, the chromatin state maps correctly group the two primary thymus tissue samples (E093 and E112) together with purified T-cells (E033-E062). We also noticed that the heart and muscle cells types are proximal to each other, which is consistent with their functional similarity. For unknown reasons, the brain cell types were divided into two distinct tight clusters. Of note, the 16 cultured cell lines from ENCODE2012 were separated from the cell types, suggesting fundamental chromatin state differences between cultured and primary cells. More complete information about the cell-type similarity can be seen in **Supp Fig. 5**.

**Figure 2.**
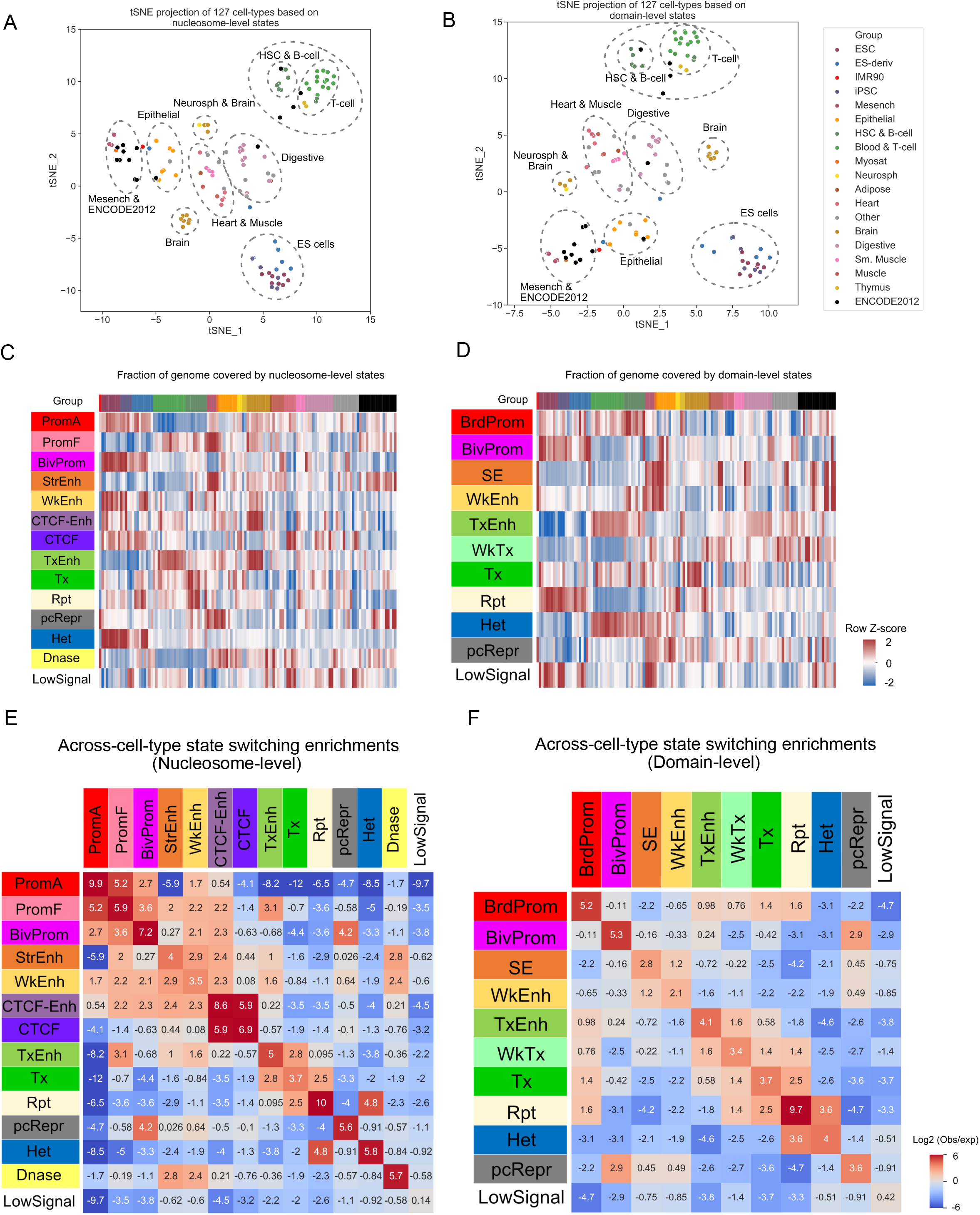
Variability analysis of the nucleosome- and domain-level chromatin states. (A-B) t-SNE projection of the distance matrix of the diHMM states between different cell types. (A) nucleosome-level (B) and domain-level. The group membership information was obtained from the Roadmap consortium. (C-D) Heatmap showing the genomic coverage for each state in each cell-type. Celltypes were ordered according to the tissue type and cell origins annotated by the Roadmap consortium. (E-F) Heatmap showing the relative frequency of chromatin state switching across cell-types. The number shown in each box represents the log2 (observed/expected) enrichments.

The availability of a consensus map in 127 cell types allowed us to systematically characterize the variation of chromatin states at both nucleosome- and domain-levels. First, we noticed that the overall coverage of each chromatin state is highly cell-type specific (Figure **2C-D**). The Active-Promoter and Bivalent-Promoter states are highly abundant in the ES cells but become less so in differentiated cells. Conversely, both Strong-Enhancer and Transcribed & Enhancer states are depleted in ES cells. This trend is consistent with our recent findings where there is a switch between promoter-centric to enhancer-centric logic with the age of ontology [11].

The overall variability associated with each chromatin state is quantified by the consistency of state presence at each genomic locus (see **Methods**, **Supp Figure 6**). As expected, both Strong Enhancer (nucleosome-level) and Super-Enhancer (domain-level) states display much higher specificity, whereas the Low-Signal state, which has the largest genome coverage, is least variable (**Supp Figure 6B and D**). Of the two CTCF associated nucleosome-level states, the CTCF-Enhancer state shows a much higher degree of cell-type specificity as compared to the CTCF state, suggesting the elements corresponding to these states may have distinct biological functions.

Next, we explored the pattern of chromatin state switching across different cell-types. (**Figure 2E-F**). We observed frequent switches between Active-Promoter and Promotor-Flanking states, reflecting promoter activity changes. Interestingly, the enhancer states (Strong, Weak, or Transcribed Enhancers) tend to transition to other states promiscuously, indicating a much higher degree of plasticity.

### Context-specific functionality of nucleosome-level chromatin states

Multi-scale chromatin state annotation provides a unique opportunity to characterize the functionality of a regulatory element in a broader context. In previous work [6], we showed that the contribution of an enhancer element to gene expression varies significantly according to the corresponding domain-level states. Here we expanded the analysis by using the annotations from all 127 cell types. Consistent with our previous findings, genes associated with Strong-Enhancer (nucleosome-level state) in the Super-Enhancer domain have higher expression levels as compared to those falling into the Weak-Enhancer domain (**Figure 3A**). Similarly, an Active-Promoter (nucleosome-level state) can be found in the Broad-Promoter domain or the Bivalent-Promoter domain. We found that the genes in the Broad-Promoter domain on average have higher expression levels than those in the Bivalent-Promoter domain. Genes associated with the Bivalent-Promoter nucleosome-level state have higher expression levels in the Bivalent-Promoter domain as compared to the Polycomb-Repressed domain (**Figure 3A**).

**Figure 3.**
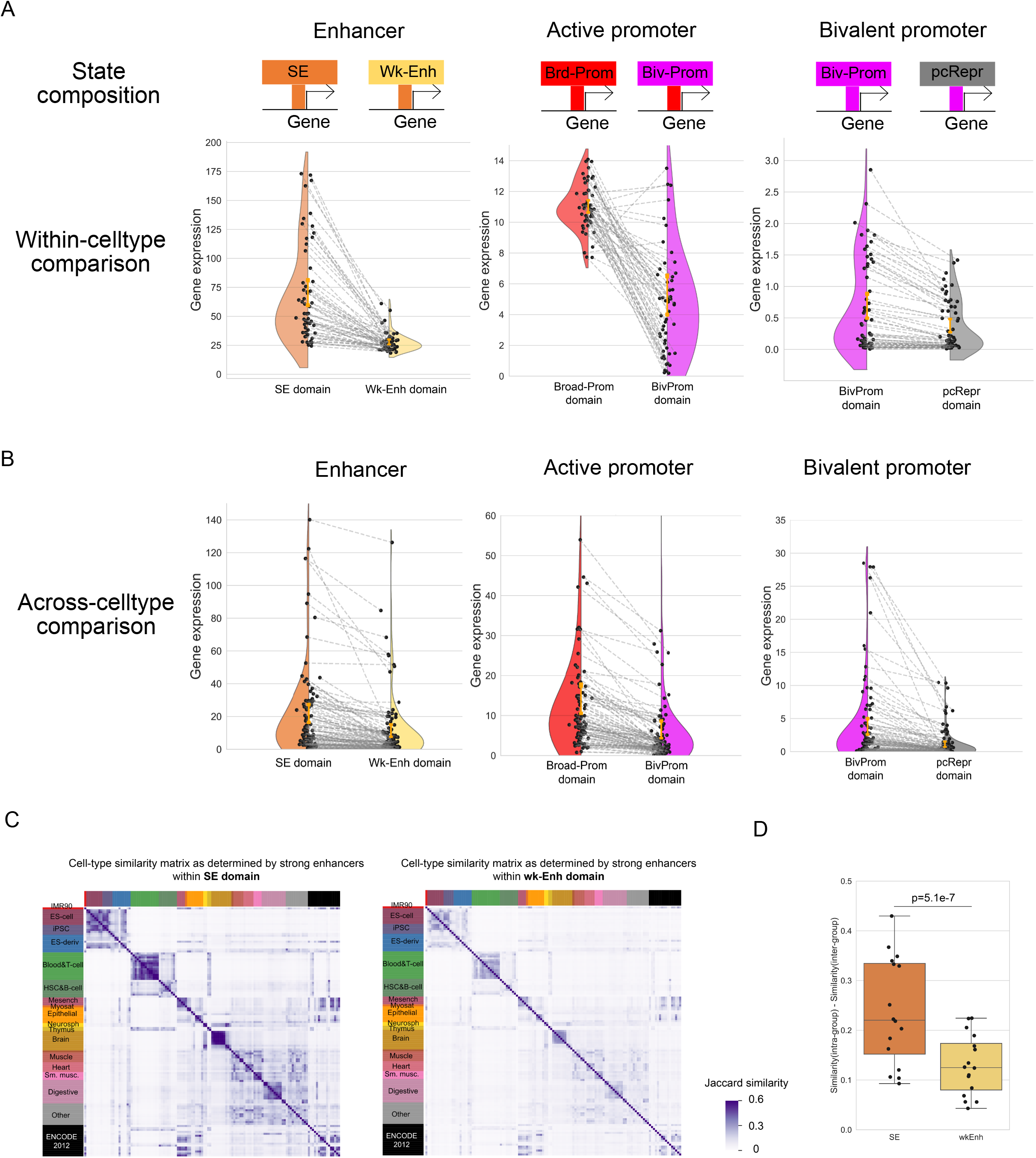
Context-specific functions of nucleosome-level states. (A) Violin plots showing the within-cell-type comparisons for genes mapped to corresponding state compositions, as shown at the top. Each dot represents the average expression of the genes that are associated with the indicated nucleosome- and domain-level states. (B) Violin plots showing the across-cell-type comparison for genes mapped to corresponding state compositions. Each dot represents the average expression of the genes that are associated with the indicated nucleosome- and domain-level states. (C) Jaccard similarity matrix showing the similarity between cell-types based on Strong-Enhancer (N8-9) in different domain-level states (D5-6, Super-Enhancer domain and D7, Weak-Enhancer domain). Cell-types were ordered according to the tissue type and cell origins annotated by the Roadmap consortium. (D) Boxplot showing the difference in revealing the cell identity between using strong enhancers in the SE or Weak-Enh domains. P-value was calculated using the Mann-Whitney U test.

Furthermore, we also tested whether the switch of chromatin states across cell types are associated with gene expression changes in a domain-specific manner. As control, we focused on cases whether the nucleosome-level states remain unchanged, but the domain-level states changed. As an example, we considered strong enhancers whose domain-level states changed from the Super-Enhancer to Weak-Enhancer domain. The gene expression levels decreased significantly (**Figure 3B**). Similarly, an Active-Promoter that switched from the Broad-Promoter to Bivalent-Promoter domain state was typically associated with reduced gene expression. These analyses strongly suggest that the domain-level chromatin state plays an important role in gene regulation.

Due to the strong cell-type specificity of super enhancers [12], we were interested to evaluate to what extent the Super-Enhancer domain distribution alone was able to distinguish different cell types. To this end, we applied the Jaccard-similarity matrix to quantify the agreement between Super-Enhancer domain distributions in different cell types (**Figure 3C, left panel**). The resulting block-diagonal pattern indicates that cells from different cell types are typically associated with distinct super-enhancer landscape. For comparison, we repeated the analysis based on the Weak-Enhancer domain distribution (**Figure 3C, right panel**). As expected, the ability to distinguish different cell types was much weaker (**Figure 3D**).

### Domain-level chromatin states reveal known and novel features associated with higher-order chromatin organization

It is known that there is a strong association between the 3D chromatin structure and the 1D chromatin state distribution. A number of computational methods have been developed to predict the 3D chromatin structure from 1D chromatin states [13–17] with various degrees of success. We recognized that many well-characterized 3D chromatin structures are associated with broad chromatin scale, therefore thought it would be natural to test whether the domain-level annotation provided by the diHMM model is useful for identifying such associations. To this end, we used enrichment analysis to identify specific associations between the domain-level chromatin states and a set of well-characterized 3D chromatin structures, including boundaries of Topologically Associating Domains (TADs), Hi-C interaction hubs (also known as Frequently Interacting Regions [18]), sub-compartments [19] and replication timing. For comparison, we repeated the same procedure to analyze the association between nucleosome-level states and various 3D chromatin features. Due to the limited availability of high-resolution Hi-C data, we only considered two cell types (E116, GM12878 lymphoblastoid cells; E017, IMR90 fetal lung fibroblasts) which high-resolution Hi-C data was publicly available.

Consistent with previous findings [20,21], we found that both nucleosome- and domain-chromatin states can be roughly divided into two groups: gene-associated (N1-N25, D1-D18) vs gene-depleted (N26-N30, D19-D30) states. While the gene-associated states are generally associated with TAD boundary, interaction hubs, and A-compartments, the gene-depleted states tend to be associated with B-compartments (**Figure 4A-B**).

**Figure 4.**
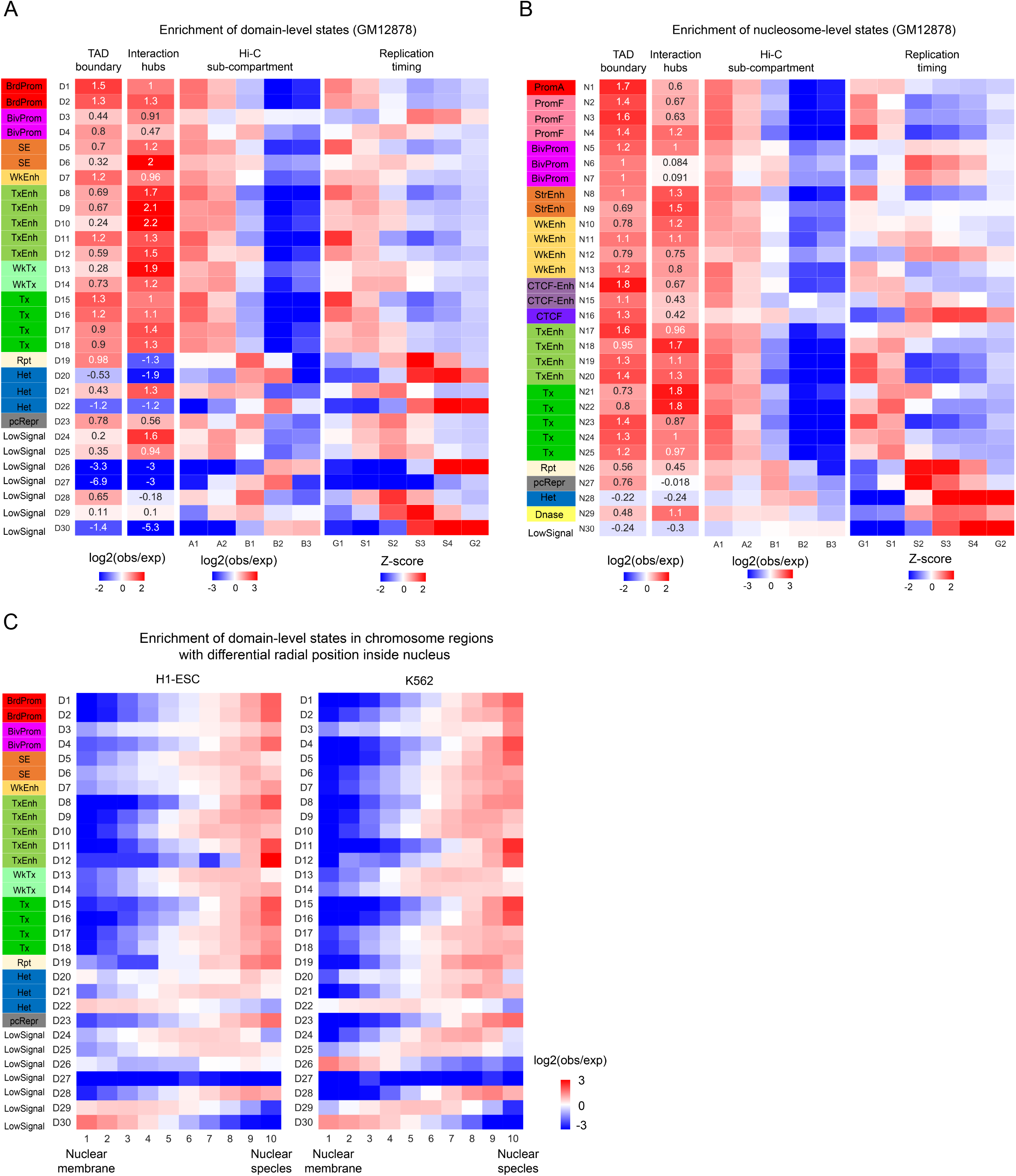
Associations between chromatin states and higher-order chromatin structures. (A-B) Enrichment analysis of domain- (A) and nucleosome-level (B) states in a set of well-characterized 3D chromatin features in GM12878 cells. (C) Enrichment analysis of domain-level states in different nuclear regions.

At the same time, we also observed certain new patterns that may be functionally relevant. For example, while the Bivalent-Promoter domain states D3 and D4 have similar histone modification patterns (see **Figure 1C**), they have distinct associations with certain 3D chromatin features. Specifically, the D3 state is strongly associated with late replication timing, while the opposite trend is associated with the D4 state (**Figure 4A** and **Supp Figure 7A**). Similarly, of the three Heterochromatin domain states (D20-D22), the D21 and D22 states have distinct association patterns with certain 3D chromatin features, with D21 enriched in A-compartments whereas D22 enriched in B-compartments. Consistent with this observation, D21 is distributed closer to nuclear speckles, whereas D22 is closer to the nuclear membrane (**Figure 4C**). Additional tests are needed in the future to help understand the implications of these observations.

## Discussion

The 3D Chromatin is hierarchically organized. In previous work [6], we developed a diHMM model to systematically characterize the chromatin states at multiple scales. However, the original study was limited by its computational inefficiency. Here, we improved the computational efficiency through optimized implementation and a consensus-clustering-based divide-and-conquer approach, which allowed us to annotate chromatin states in 127 human cell types using the data obtained from the Epigenome Roadmap and ENCODE consortia.

This multi-scale chromatin state map allowed us to systematically investigate the effect of spatial context in mediating the functionality of a regulatory element. For several nucleosome-level states (Strong-Enhancer, Active-Promoter, and Bivalent-Promoter), we demonstrated that its role in gene regulation is dependent on the domain-level context. Thus, our multi-scale chromatin state annotation has provided new insights into the mechanism of gene regulation. In addition, we also identified previously unrecognized relationship between 1D chromatin states and 3D chromatin features such as the distinct association between two Bivalent-Promoter domain states with replication timing. Taken together, our analyses have demonstrated the utility of our multi-scale chromatin state map as a valuable resource for the community.

## Methods

### Data sources

We performed the diHMM annotation based on the imputed binarized signals at 200 bp resolution, including 11 histone marks (H3K4me3, H3K4me2, H3K4me1, H3K27ac, H3K9ac, H2A.Z, H3K79me2, H4K20me1, H3K36me3, H3K9me3, H3K27me3), DNase hypersensitivity and CTCF. The full 12-mark epigenome signal matrix of the 127 cell-types, which are at the resolution of 200bp, were downloaded from [https://personal.broadinstitute.org/jernst/BINARY_IMPUTED12/]. CTCF signals were obtained by using imputations as described in the section below. We used the imputed signal here because not every epigenomic mark is commonly available in all the cell-types. Detailed information of the 127 cell-types (epigenomes) can be found at the Roadmap website (http://www.roadmapepigenomics.org).

### Imputation of CTCF tracks

The signal track of CTCF for different cell types was imputed by ChromImpute [22]. Briefly, the processing consisted of the following steps: (1) Processed files which contain signal p-values for CTCF were downloaded from ENCODE. In total, CTCF tracks for 10 cell-types (E114, E116, E117, E118, E119, E121, E123, E124, E125 and E127) were obtained. Then the “Convert” function from ChromImpute [22] was applied to convert the data into desired resolution (25bp). (2) The processed CTCF signals were combined with the full matrix with observed epigenomic signals from ROADMAP. (3) “ComputeGlobalDist”, “GenerateTrainData”, “Train” and “Apply” functions from ChromImpute software were sequentially used to finally impute signal tracks for all cell types. (4) Binarized CTCF signal tracks with threshold of 2 based on the average of the eight 25bp bins in the 200bp interval.

### Training diHMM models for each cell-type

The diHMM models for each cell-type were trained using the C++ diHMM package, with the numbers of nucleosome- and domain-level states set to 30. The bin sizes for nucleosome- and domain-level states were set to 200bp and 4 kb respectively. 1e-6 was used for the tolerance value for the model to converge, and 500 was set to the maximum iteration.

We compared the performance of the C++ and MATLAB version by using the concatenated chr17 in GM12878 (E116), H1-ESC (E003) and K562 (E123), with Linux Ubuntu (Version 16.04.6 LTS), 12 CPUs (Intel(R) Xeon(R) CPU E5-2699 v3 @ 2.30GHz) and Mem 1T. The MATLAB implementation takes ~10 days, while the C++ version takes ~7h, which is ~30 times faster. Training a diHMM model using the C++ implementation for one cell-type usually takes ~3 days using the improved C++ diHMM algorithm, while it is infeasible using the MATLAB implementation.

### Joint segmentation in 127 cell-types using a consensus clustering approach

To generate the consensus chromatin state labels for each cell-type and obtain a unified diHMM model, we first applied the 127 models trained from 127 individual cell-types to each epigenome, resulting in 127 sets of chromatin state labels for each celltype at both the nucleosome- and domain-level. For each set of chromatin state labels at one length-scale, the chromatin states from different chromosomes were concatenated, yielding a long 1-dimensional vector for each cell-type with integer numbers ranging from 1 to 30 representing 30 different chromatin states. Each cell-type has 127 such chromatin state labels at nucleosome- and domain-level. We then adopted a modified Meta-CLustering Algorithm (MCLA), an efficient consensus clustering method [7], to obtain a set of consensus chromatin state labels for the two length-scales respectively. Specifically, consensus chromatin states were obtained by following steps:

#### Constructing chromatin state indicator vectors

State indicator vectors are binary vectors to represent the positions for one chromatin state label in the genome. The dimension of the vector is the same as the number of bins in the genome, where 1 represents that the underlying genomic bin is labeled as that chromatin state. Each set of chromatin state labeling will have 30 state indicator vectors. In total, this step led to 3810 state indicator vectors.

#### Clustering chromatin state indictor vectors

To reveal the degree of similarity among the 3810 states, we calculated the overlap between each pair of state labels by using the Jaccard index, which is defined as the ratio of the intersection to the union of a pair of state indicator vectors, between the state indicator vectors, resulting in a Jaccard similarity matrix with a dimension of 3810*3810.

#### Collapsing state indicator vectors into a selected number of meta-states

To find matching states labels in the 3810 states, we performed hierarchical clustering to the Jaccard similarity matrix obtained above and cut the dendrogram at a distance which would result in 30 meta-clusters. Each meta-cluster consists of multiple chromatin state labels from different models that are highly overlapped with each other. The 127 sets of chromatin state labels were then rewritten by the meta-clusters.

#### Competing for state labels for meta-states

In this step, each bin is assigned to its most associated meta-cluster across the 127 state labels by the majority vote rule. In case of ties, the bin will be attributed to a random meta-cluster with the highest association.

### Assessing the degree of shared information between chromatin state maps resulting from different models

To measure the agreement of chromatin state labels on the same epigenome resulting from models trained from different cell-types, we used Normalized Mutual Information (NMI), a metric for quantifying the statistical information shared between two distributions. NMI is a bounded value ranging from 0 (no shared information) to 1 (perfect matching). The function “mutual_info_score” from the scikit-learn package [23] was used for the calculation. Specifically, we first applied the diHMM models trained from each celltype to the epigenome of all cell-types and obtained the consensus labels, resulting in 127 (individual cell-type) and 1 (consensus) chromatin state labels for each cell-type. Then NMI was calculated for each pair of the chromatin state labels for each cell-type, giving a symmetric NMI matrix with a 128*128 dimension. Then we obtained the average NMI matrix from the 127 cell-types. The mean NMI between one cell-type and all other cell-types was calculated and plotted in **Sup Figure 2**.

### Mnemonics assignment of nucleosome- and domain-level chromatin states

The nucleosome-level chromatin state labels were interpreted into functional terms by using the combinatorial pattern of chromatin marks, enrichment on functional elements and genomic coverage. The combinatorial pattern of chromatin marks associated with each state were represented by the empirical emission matrix, where each row shows the probabilities that each chromatin mark was found in the state. These probabilities were calculated after concatenating the consensus chromatin state labels of the 127 cell-types. For the chromatin state enrichment on functional elements shown in **Supp Figure 3A**, we used the “annotatePeaks.pl” script from the Homer software [24] to obtain the enrichment fold change. We further normalized the enrichment pattern by the maximum value in each row.

The domain-level chromatin state labels were interpreted into functional terms by using the enrichment into the nucleosome-level chromatin states. Specifically, for each domain state, the enrichment of nucleosome-level states was calculated as the observed frequency of nucleosome-level states contained in the domain state relative to the expected frequency of states based on state coverage.

### Comparing diHMM’s nucleosome-level states with chromHMM’s annotation

To see if the nucleosome-level states are consistent with chromHMM’s annotations, we computed the similarity between our 30 states and chromHMM’s 25 states using Jaccard similarity index.

### t-SNE projection of 127 cell-types based on chromatin states

To visualize the similarity of the nucleosome- and domain-level chromatin states of the 127 cell-types, we performed hierarchical clustering of the chromatin state similarity matrix and projected the 127 cell-types in the t-SNE space. Briefly, we first calculated the similarity between the chromatin states of any two epigenomes using NMI, which gave a 127*127 similarity matrix. Since a large portion of the genome is quiescent, we filtered out those regions where all the 127 cell types have the “LowSignal” states. Then we obtained the distance matrix of epigenomes by take 1 – NMI similarity matrix. The distance matrix was then used as the precomputed matrix to the t-SNE function in scikit-learn [23] and the coordinates of each cell-type in the t-SNE space were calculated.

### Chromatin state switching between cell-types

The fold enrichments represent the relative frequency that a genomic position annotated with one chromatin state in one cell-type will switch to the indicated chromatin state in a second cell-type, after controlling the state coverage. Values are relative enrichments calculated in aggregating over all possible pairwise comparisons of cell-types.

### Context-specific functions of nucleosome-level chromatin states

To demonstrate that nucleosome-level chromatin states’ functions are dependent on the domain state, we compared the expressions of genes associated with the same type of nucleosome-level states but different domain types. Using the active promoter in broad promoter and bivalent promoter domains as an example, we performed the within- and across-cell-type comparisons as follows.

For within-cell-type comparison, we first selected the nucleosome-level “active promoter” state (N1) locating within “Broad promoter” (D1 and D2) or “Bivalent promoter” (D3 and D4) domains respectively. Then the two sets of active promoters were associated with nearest genes, and only associated genes with genomic distance shorter than 4kb were kept. The two sets of associated genes were compared and the genes specific to each set were used for comparison.

For across-cell-type comparison, we focused on genes that are associated with same nucleosome-level states but different domain types. For each gene, we divided cell-types into two groups: the cell-types in which the promoter of the gene was associated with active promoter state (N1) in the broad promoter domain (D1 or D2), and the cell-types in which the promoter of the gene was associated with nucleosome-level active promoter state and bivalent promoter domain state (D3 or D4). The mean expression level of the two groups of cell-types were compared for each gene. Only genes with more than 5 cell-types in each group were kept for analysis.

Same approach was used for the comparison of bivalent promoter in the bivalent promoter and Polycomb-repressed domains. For comparing the enhancer in the SE and Wk-enh domains, we modified the gene association rule as the following: only genes with genomic distance shorter than 10kb were kept.

### Chromatin state enrichment in higher-order genome organization features

We tested the association between the multi-length-scale chromatin states and the higher-order genome organization features, including TAD boundaries, Hi-C interaction hubs, sub-compartments and replication timing. We chose two cell-types (GM12878 and IMR90) which have the most comprehensive 3D genome features to demonstrate the association. Specifically, TADs were identified from [19] as identified by the Arrowhead algorithm at the resolution of 40kb. The first and last bin of TADs were extended by 40kb at both directions and were used as TAD boundaries. The enrichment of a particular chromatin state in TAD boundaries was calculated as 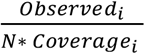, where *Observed_i_* is the observed frequency of state *i* intersecting with TAD boundaries, *N* is the total number of chromatin states of TAD boundaries and *Coverage_i_* is the genome-wide fraction of state *i*. Hi-C interaction hubs at 40kb resolution, which are also known as Frequently interacting Regions (FIREs), were downloaded from [18]. Sub-compartments for GM12878 and IMR90 were downloaded from [25]. The enrichment of chromatin states in interaction hubs and sub-compartments were calculated in the same way as TAD boundaries.

For replication timing signals, we downloaded the bigwig tracks of the six cell-cycle phases (from early to late: G1,S1,S2,S3,S4,G2) from the ENCODE consortium and their accession numbers were summarized in **Sup Table 1**. For each chromatin state, the mean quantitative signals in each phase were calculated and normalized across the six phases using Z-score.

To validate the differences between chromatin states in their radial positions inside the nucleus, we tested the distribution of chromatin states in terms of the TSA-seq signals [26]. TSA-seq is a sequencing-based technique to estimate the mean chromosomal distance from nuclear speckles genome-wide. To see if there is a difference in the radial location of the two states, we divided the whole genome into 10 equal deciles according to their TSA-seq signals and tested the enrichment of the chromatin states in each decile. TSA-seq are available for two cell-types: H1-ESC and K562, at the resolution of 20kb. The bigwig tracks of TSA-seq signals for the two cell-types were downloaded from the 4DN project [27] with accession numbers summarized in **Sup Table 1**. First, the 20kb genomic bins were sorted by the TSA-seq signals and divided into 10 deciles of equal size. Then the obs/exp enrichment of chromatin states in each decile were calculated as described above.

### Data availability

The nucleosome- and domain-level chromatin state annotations in the 127 cell types as well as the code for implementing the consensus diHMM model are freely available to the public and can be downloaded from (https://github.com/ykai16/diHMM).

## Competing interests

The authors declare no competing interests.

## Author contributions

GCY conceived the project. ST rewrote the improved diHMM package in C++. SBS generated the CTCF imputation data. YK implemented the consensus clustering method and performed all the analyses. YK and GCY wrote the manuscript with input from all authors. GCY supervised the study.

## Acknowledgements

We thank the Roadmap and ENCODE consortia for generating the public datasets. We thank Drs. Jason Ernst, Wouter Meuleman for helpful discussions. Dr. Ernst also generously provided the imputed Epigenome Roadmap data which was analyzed here. We thank Dr. Rui Dong for helping with setting up the computing environment. We also thank the Research Computing team in Dana-Farber Cancer Institute for support with the model training. This work was supported by National Institutes of Health grants (R01HL119099 and R01HG009663) to GCY.

## Supplementary Figure Legend

**Supplementary Figure 1.**
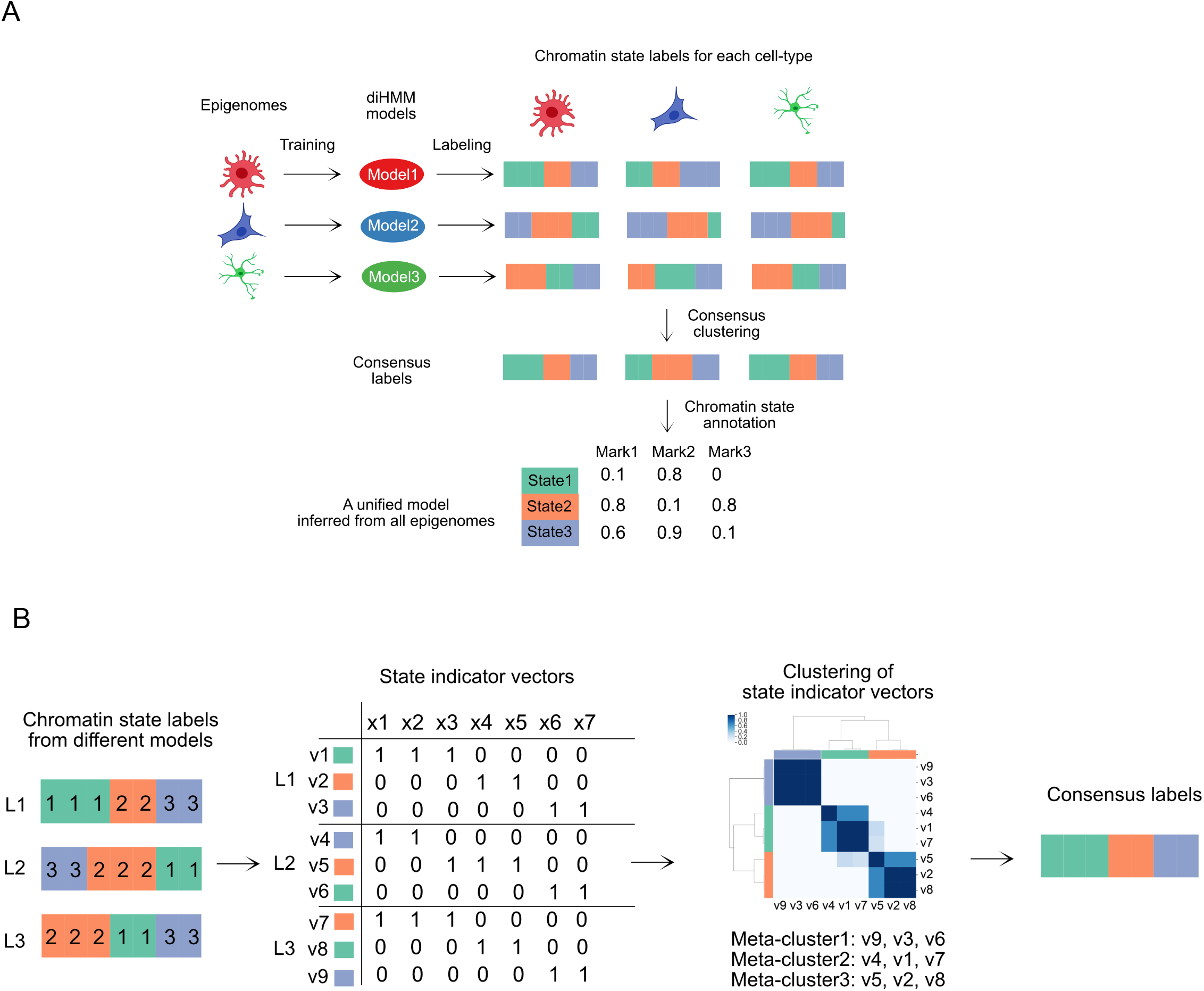
Illustration of the approach to obtain a unified annotation across 127 epigenomes. (A) Illustration of the overall approach. Four steps were involved: (1) Training a diHMM model from each epigenome; (2) Applying each diHMM model to annotate chromatin states for all the epigenomes, resulting in 127 sets of chromatin states for each epigenome; (3) Performing consensus clustering of the 127 sets of chromatin states for the concatenated epigenomes; (4) Manually annotating chromatin states based on empirical emission matrix for nucleosome-level states. (B) Illustration of the consensus clustering method for obtaining the consensus annotation from a set of cell-type-specific annotations. Three steps were involved in this method: (1) Constructing chromatin state indicator vectors. (2) Clustering chromatin state indictor vectors. (3) Collapsing state indicator vectors into a selected number of meta-states. (4) Competing for state labels for meta-states. See Methods for detailed information.

**Supplementary Figure 2.**
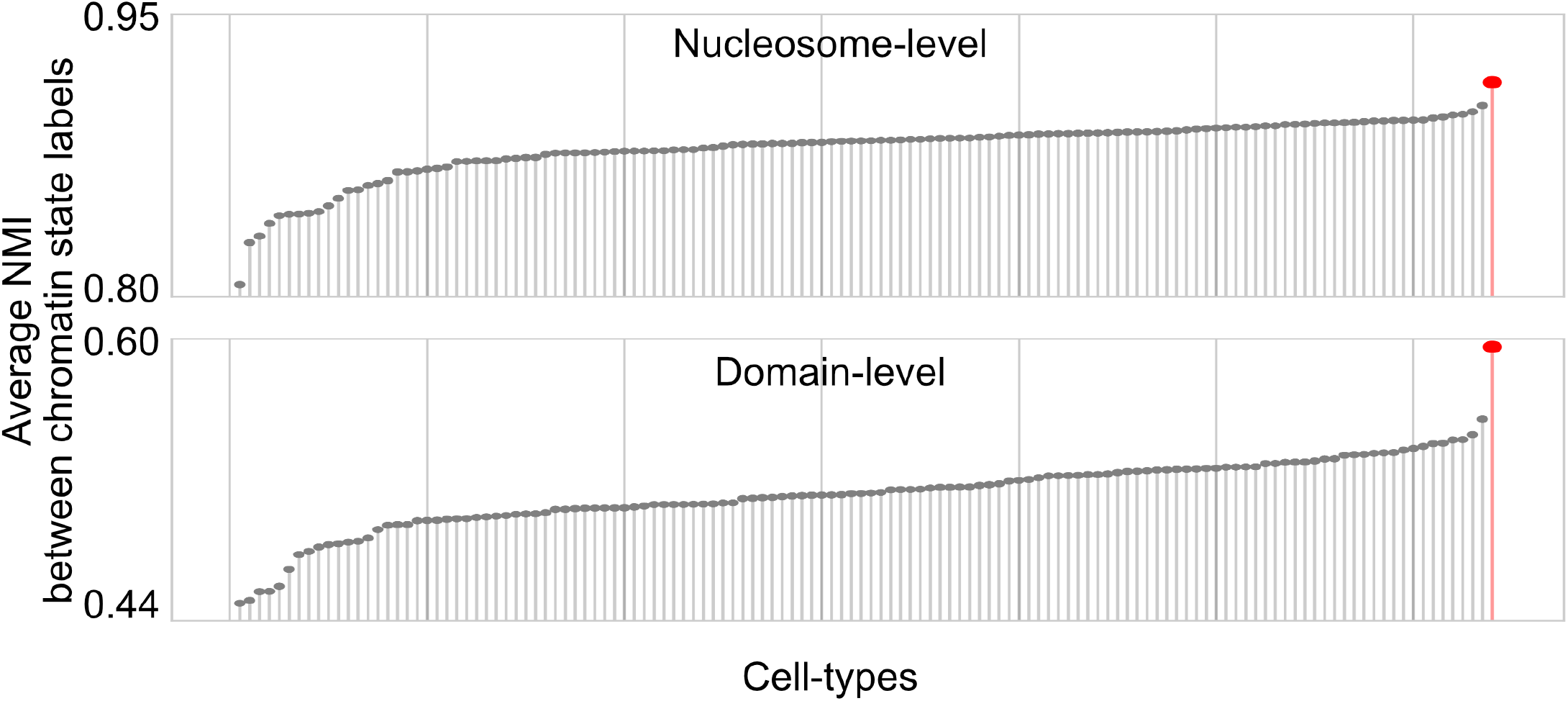
Lollipop plot showing the degree of agreement among celltype-specific annotations and consensus annotation. Each gray dot represents the average NMI (Normalized Mutual Information) between one annotation derived from one cell-type-specific model and annotations derived from models trained from other cell-types. Red dot represents the average mutual information between the consensus annotation and all the cell-type-specific annotations. Cell-types were ordered from low to high for nucleosome- and domain-level comparisons separately.

**Supplementary Figure 3.**
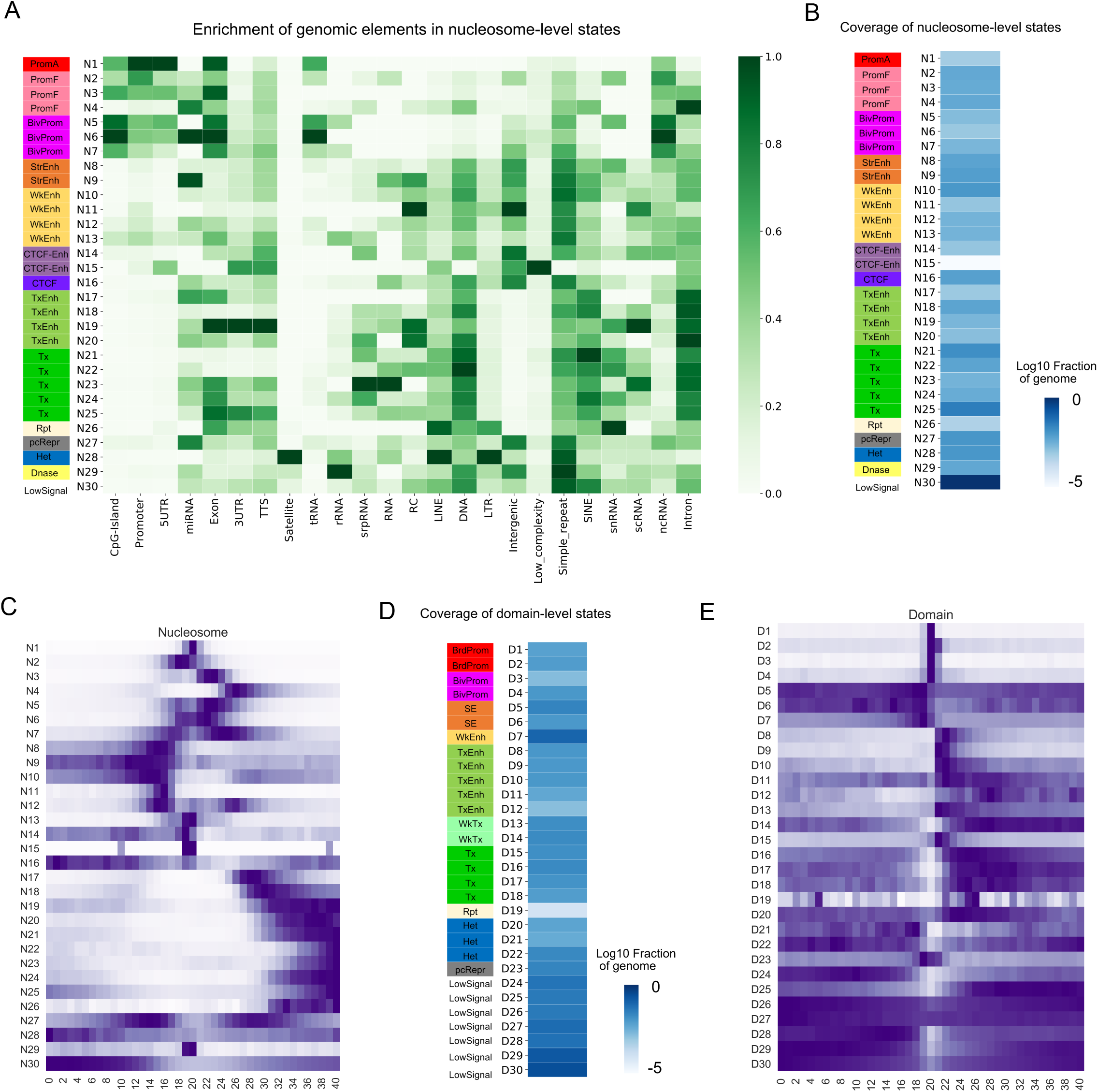
Analysis of two-length-scale chromatin states (A) Enrichment of the 30 nucleosome-level states in functional genomic elements. Data shown were from GM12878 cells. Each column shows relative enrichment in a linear scale between 0 and 1. (B) Fraction of genome coverage for the 30 nucleosome-level states in the concatenated 127 genomes. Color bar shows the log10 scale of fractions of the genome covered by each state. (C) Spatial distribution of nucleosome-level chromatin states relative to TSSs. The frequency of each state spanning over the region (TSS +/− 20 bins) was counted and normalized based on the maximum frequency at each row. (D) Fraction of genome coverage for the 30 domain-level states. (E) Spatial distribution of domain-level chromatin states relative to TSSs. The frequency of each state spanning over the region (TSS +/− 20 bins) was counted and normalized based on the maximum frequency at each row.

**Supplementary Figure 4.**
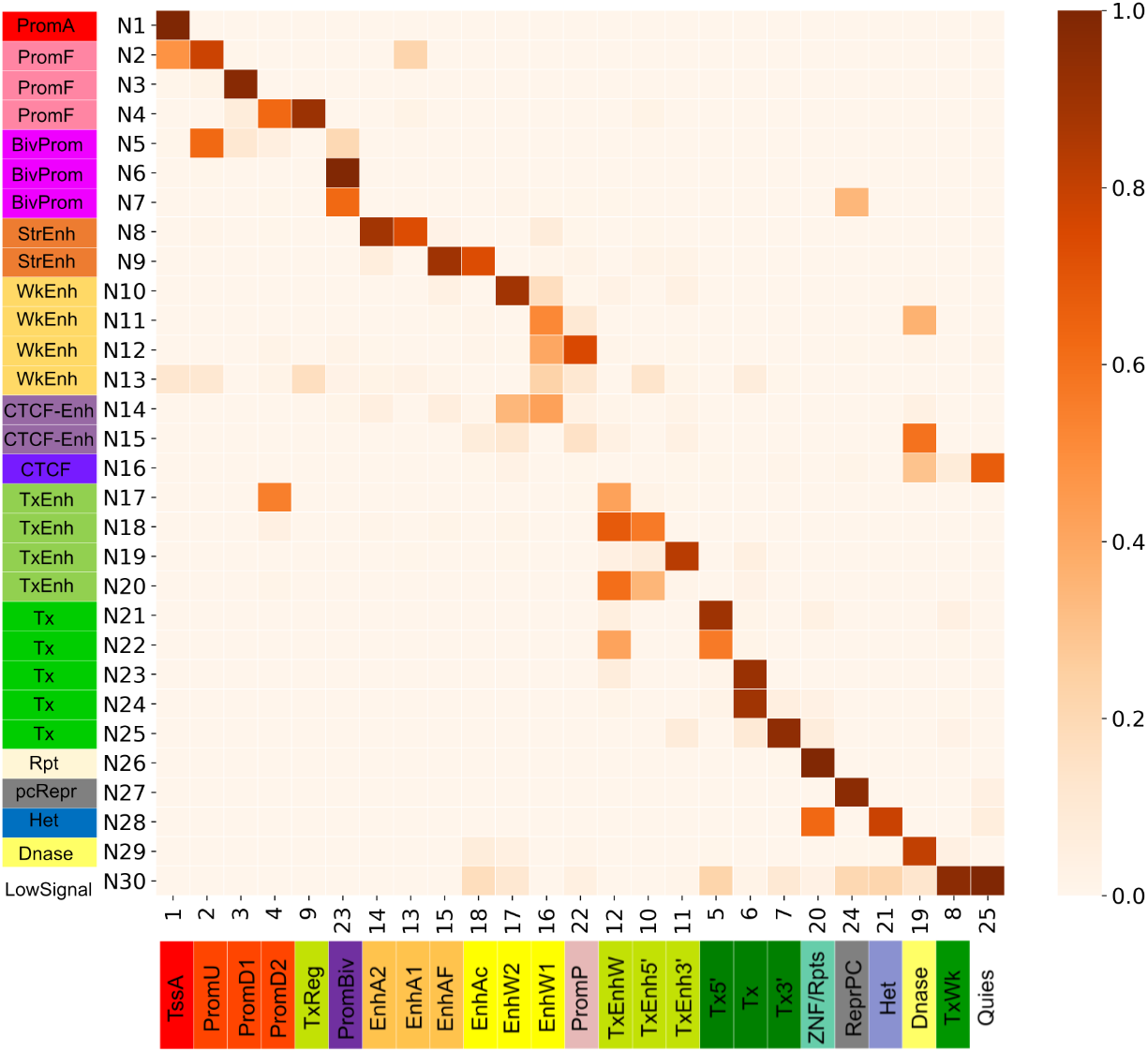
Heatmap showing the consistency between the diHMM’s nucleosome-level states and chromHMM’s annotation. Each pixel represents the Jaccard similarity between two states. GM12878 data were shown as an example.

**Supplementary Figure 5.**
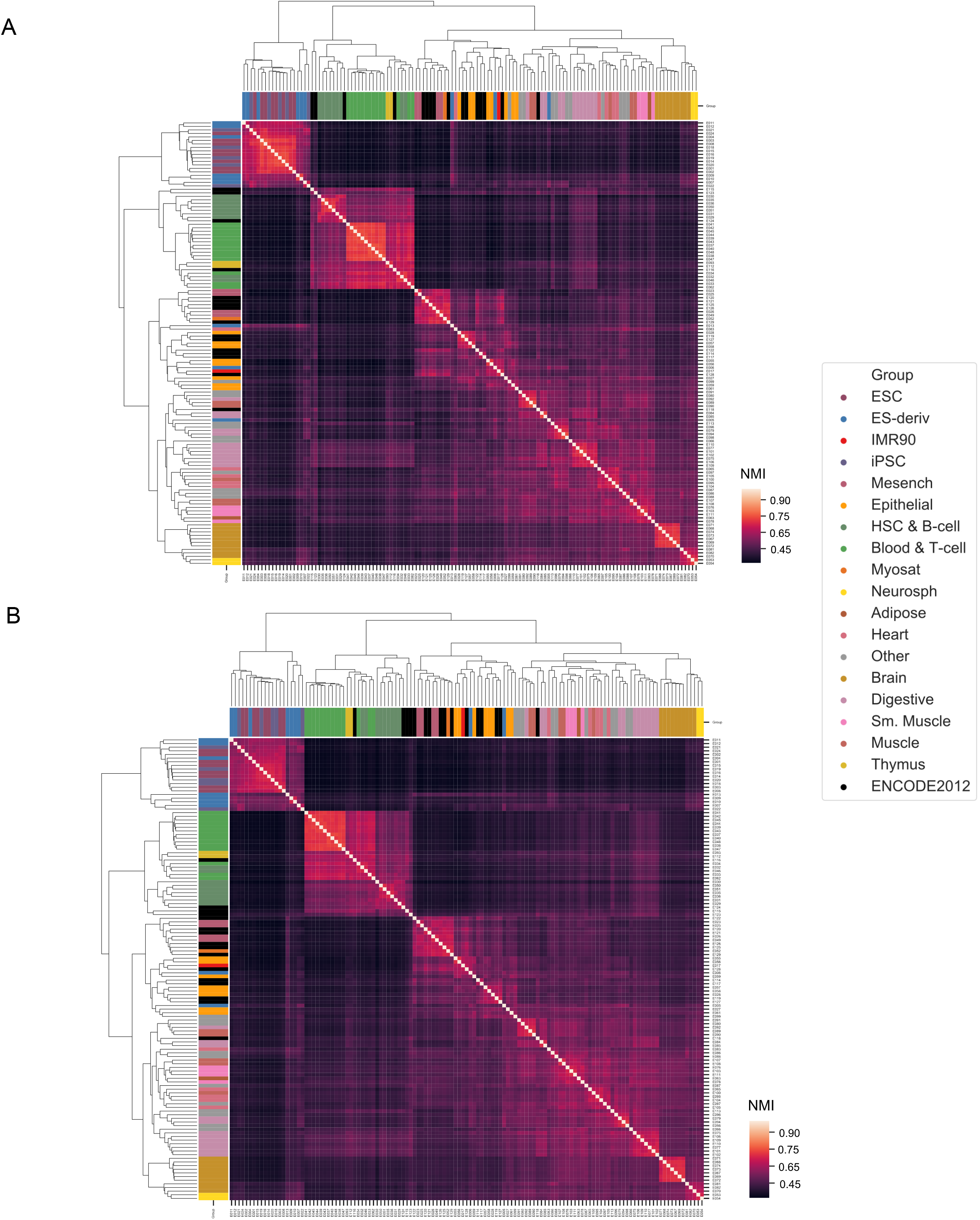
Hierarchical clustering of the similarity matrix of genome-wide nucleosome- (A) and domain-level (B) chromatin state maps across 127 epigenomes. Chromatin state similarity was calculated using NMI based on genome-wide chromatin state maps. Group membership of each epigenome was from Roadmaps. The names of epigenome could be seen by zooming in.

**Supplementary Figure 6.**
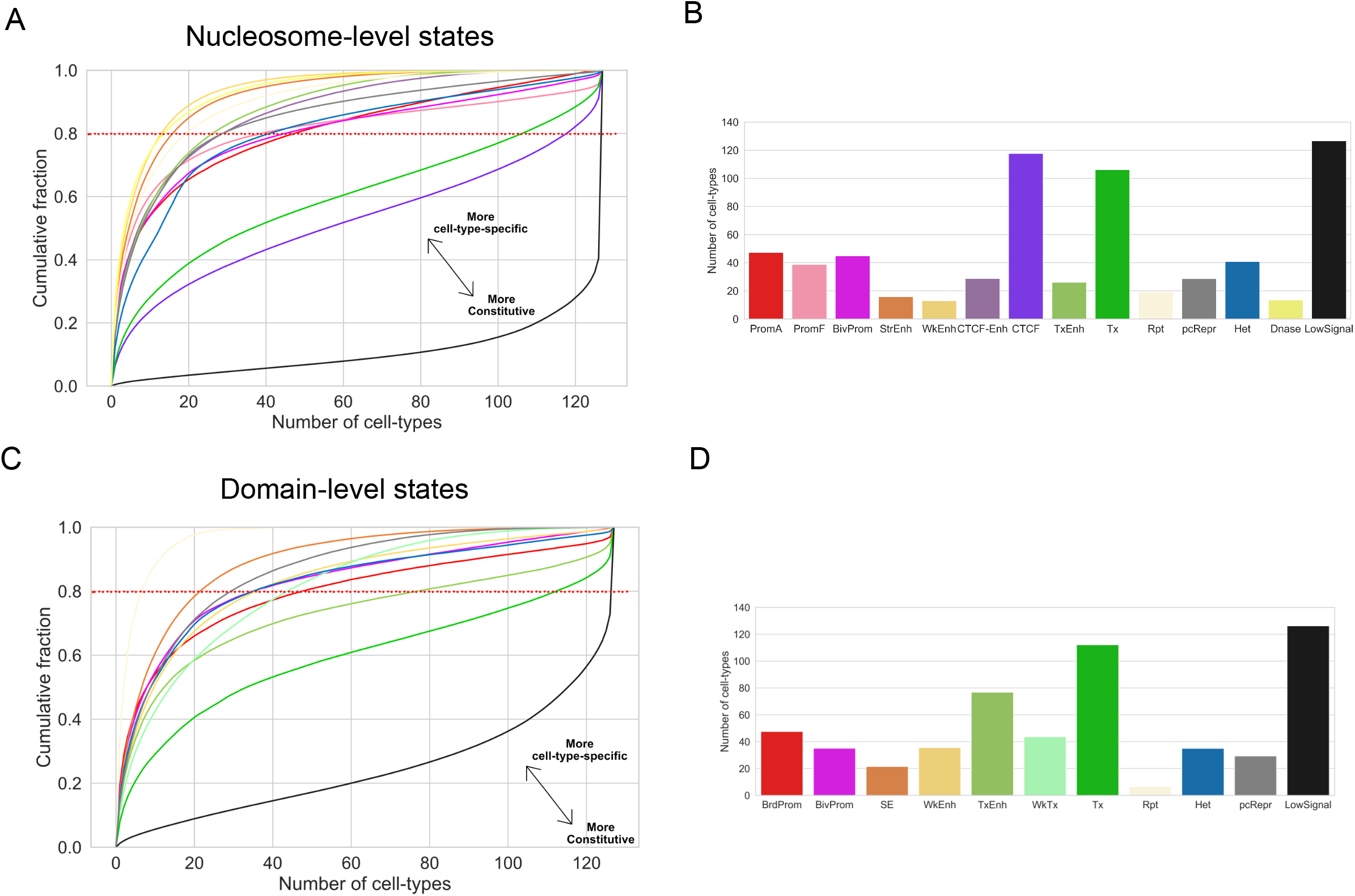
Variability of two-length-scale chromatin states. (A-C) Cumulative distribution showing variability of nucleosome- (A) and domain-level (C) states. Points in each curve represent how much fraction of the state (y-coordinate) are conserved in a certain number of epigenomes (x-coordinate). (B) Number of epigenomes covered by 80% of nucleosome- (B) and domain-level (D) state instances in the genome. The higher the number is, the more specific for the state.

**Supplementary Figure 7.**
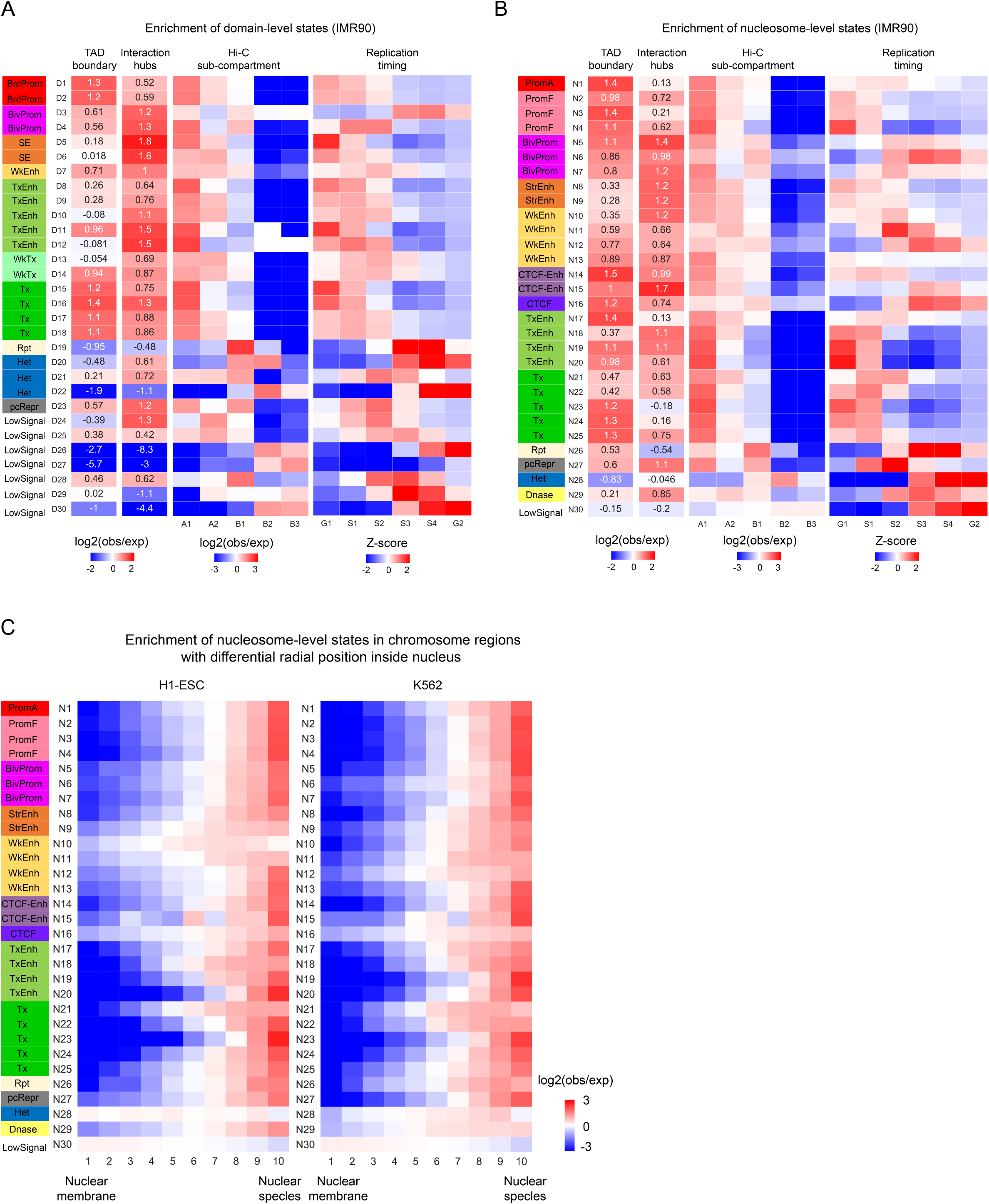
Associations between chromatin states and higher-order chromatin structures. (A-B) Enrichment of domain-level (A) and nucleosome-level (B) states in a set of features related to higher-order chromatin architecture, including TAD boundaries, Hi-C interaction hubs, Hi-C sub-compartments and replication timing. Data shown are from IMR90 cells. (C) Enrichment of nucleosome-level states in genomic regions characterized with different radial position inside nucleus.

---

**Supplementary Table 1.**
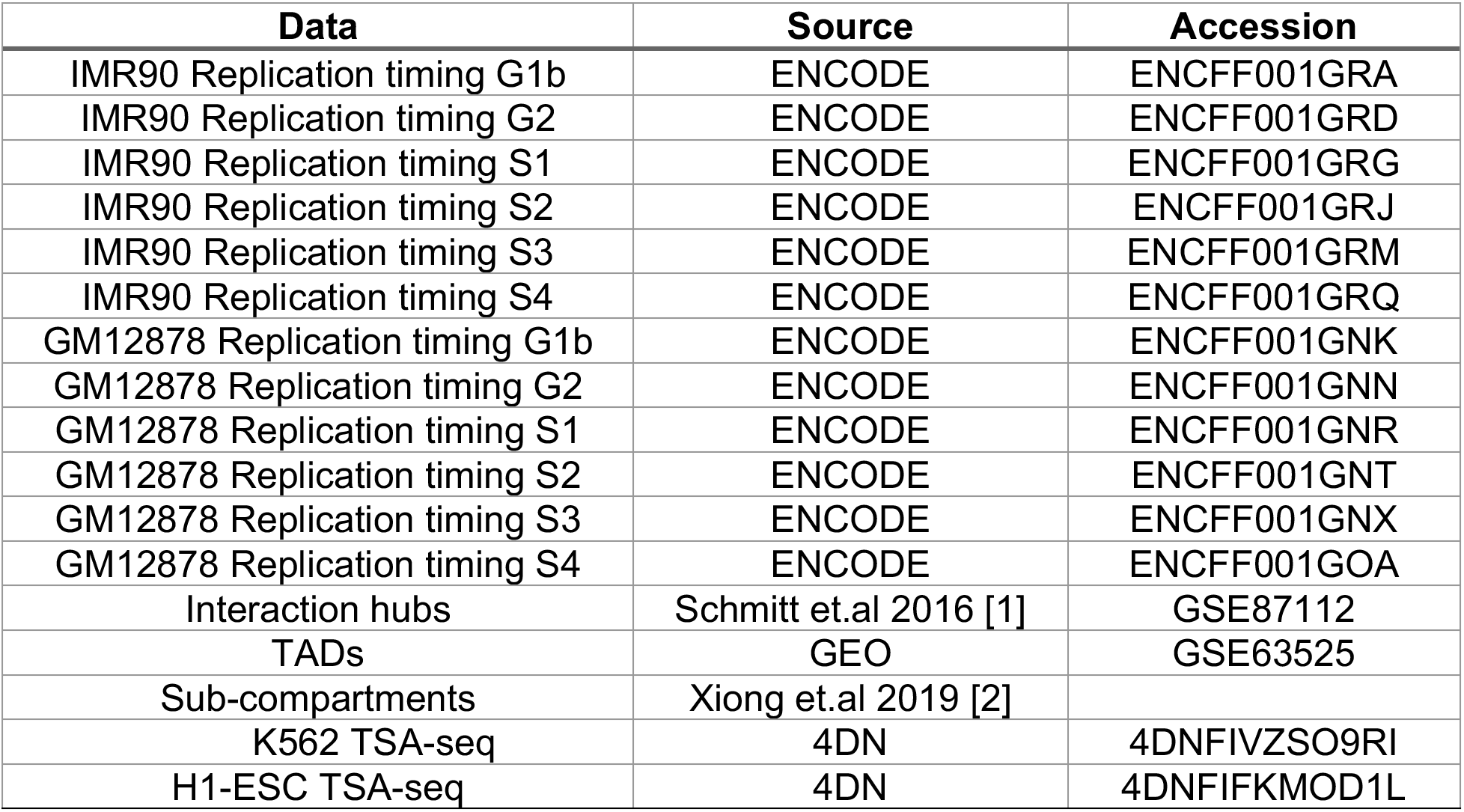
Public data used in the study

| **Data** | **Source** | **Accession** |
| --- | --- | --- |
| IMR90 Replication timing G1b | ENCODE | ENCFF001GRA |
| IMR90 Replication timing G2 | ENCODE | ENCFF001GRD |
| IMR90 Replication timing S1 | ENCODE | ENCFF001GRG |
| IMR90 Replication timing S2 | ENCODE | ENCFF001GRJ |
| IMR90 Replication timing S3 | ENCODE | ENCFF001GRM |
| IMR90 Replication timing S4 | ENCODE | ENCFF001GRQ |
| GM12878 Replication timing G1b | ENCODE | ENCFF001GNK |
| GM12878 Replication timing G2 | ENCODE | ENCFF001GNN |
| GM12878 Replication timing S1 | ENCODE | ENCFF001GNR |
| GM12878 Replication timing S2 | ENCODE | ENCFF001GNT |
| GM12878 Replication timing S3 | ENCODE | ENCFF001GNX |
| GM12878 Replication timing S4 | ENCODE | ENCFF001GOA |
| Interaction hubs | Schmitt et.al 2016 [1] | GSE87112 |
| TADs | GEO | GSE63525 |
| Sub-compartments | Xiong et.al 2019 [2] |  |
| K562 TSA-seq | 4DN | 4DNFIVZSO9RI |
| H1-ESC TSA-seq | 4DN | 4DNFIFKMOD1L |

## Notes

https://github.com/ykai16/diHMM

### Competing Interest Statement

The authors have declared no competing interest.

## References

1. Kouzarides T. Chromatin modifications and their function. Cell. 2007;128:693–705.

2. Ernst J, Kellis M. ChromHMM: automating chromatin-state discovery and characterization. Nat Methods. 2012;9:215–6.

3. Hoffman MM, Buske OJ, Wang J, Weng Z, Bilmes JA, Noble WS. Unsupervised pattern discovery in human chromatin structure through genomic segmentation. Nat Methods. 2012;9:473–6.

4. Filion GJ, van Bemmel JG, Braunschweig U, Talhout W, Kind J, Ward LD, et al. Systematic protein location mapping reveals five principal chromatin types in Drosophila cells. Cell. 2010;143:212–24.

5. Kharchenko PV, Alekseyenko AA, Schwartz YB, Minoda A, Riddle NC, Ernst J, et al. Comprehensive analysis of the chromatin landscape in Drosophila melanogaster. Nature. 2011;471:480–5.

6. Marco E, Meuleman W, Huang J, Glass K, Pinello L, Wang J, et al. Multi-scale chromatin state annotation using a hierarchical hidden Markov model. Nat Commun. 2017;8:15011.

7. Strehl A, Ghosh J. Cluster Ensembles --- A Knowledge Reuse Framework for Combining Multiple Partitions. J Mach Learn Res. 2002;3:583–617.

8. Consortium RE, Others. Anshul Kundaje, Wouter Meuleman, Jason Ernst, Misha Bilenky, Angela Yen, Alireza Heravi-Moussavi, Pouya Kheradpour, Zhizhuo Zhang, Jianrong Wang, et al. Integrative analysis of 111 reference human epigenomes. Nature. 2015;518:317–30.

9. ENCODE Project Consortium. An integrated encyclopedia of DNA elements in the human genome. Nature. 2012;489:57–74.

10. Maaten L van der, Hinton G. Visualizing Data using t-SNE. J Mach Learn Res. 2008;9:2579–605.

11. Cai W, Huang J, Zhu Q, Li BE, Seruggia D, Zhou P, et al. Enhancer dependence of cell-type-specific gene expression increases with developmental age. Proc Natl Acad Sci U S A. 2020;117:21450–8.

12. Hnisz D, Abraham BJ, Lee TI, Lau A, Saint-André V, Sigova AA, et al. Super-Enhancers in the Control of Cell Identity and Disease [Internet]. Cell. 2013. p. 934–47. Available from: http://dx.doi.org/10.1016/j.cell.2013.09.053

13. Huang J, Marco E, Pinello L, Yuan G-C. Predicting chromatin organization using histone marks. Genome Biol. 2015;16:162.

14. Zhu Y, Chen Z, Zhang K, Wang M, Medovoy D, Whitaker JW, et al. Constructing 3D interaction maps from 1D epigenomes. Nat Commun. 2016;7:10812.

15. Kai Y, Andricovich J, Zeng Z, Zhu J, Tzatsos A, Peng W. Predicting CTCF-mediated chromatin interactions by integrating genomic and epigenomic features. Nat Commun. 2018;9:4221.

16. Li W, Wong WH, Jiang R. DeepTACT: predicting 3D chromatin contacts via bootstrapping deep learning. Nucleic Acids Res. 2019;47:e60.

17. Qi Y, Zhang B. Predicting three-dimensional genome organization with chromatin states. PLoS Comput Biol. 2019;15:e1007024.

18. Schmitt AD, Hu M, Jung I, Xu Z, Qiu Y, Tan CL, et al. A Compendium of Chromatin Contact Maps Reveals Spatially Active Regions in the Human Genome. Cell Rep. 2016;17:2042–59.

19. Rao SSP, Huntley MH, Durand NC, Stamenova EK, Bochkov ID, Robinson JT, et al. A 3D Map of the Human Genome at Kilobase Resolution Reveals Principles of Chromatin Looping [Internet]. Cell. 2014. p. 1665–80. Available from: http://dx.doi.org/10.1016/j.cell.2014.11.021

20. Lieberman-Aiden E, van Berkum NL, Williams L, Imakaev M, Ragoczy T, Telling A, et al. Comprehensive mapping of long-range interactions reveals folding principles of the human genome. Science. 2009;326:289–93.

21. Dixon JR, Selvaraj S, Yue F, Kim A, Li Y, Shen Y, et al. Topological domains in mammalian genomes identified by analysis of chromatin interactions. Nature. 2012;485:376–80.

22. Ernst J, Kellis M. Large-scale imputation of epigenomic datasets for systematic annotation of diverse human tissues. Nat Biotechnol. 2015;33:364–76.

23. Pedregosa F, Varoquaux G, Gramfort A, Michel V, Thirion B, Grisel O, et al. Scikit-learn: Machine learning in Python. the Journal of machine Learning research. JMLR.org; 2011;12:2825–30.

24. Heinz S, Benner C, Spann N, Bertolino E, Lin YC, Laslo P, et al. Simple combinations of lineage-determining transcription factors prime cis-regulatory elements required for macrophage and B cell identities. Mol Cell. 2010;38:576–89.

25. Xiong K, Ma J. Revealing Hi-C subcompartments by imputing inter-chromosomal chromatin interactions. Nat Commun. 2019;10:5069.

26. Chen Y, Zhang Y, Wang Y, Zhang L, Brinkman EK, Adam SA, et al. Mapping 3D genome organization relative to nuclear compartments using TSA-Seq as a cytological ruler. J Cell Biol. 2018;217:4025–48.

27. Dekker J, Belmont AS, Guttman M, Leshyk VO, Lis JT, Lomvardas S, et al. The 4D nucleome project. Nature. 2017;549:219–26.

## References

1. Schmitt AD, Hu M, Jung I, Xu Z, Qiu Y, Tan CL, Li Y, Lin S, Lin Y, Barr CL, Ren B: A Compendium of Chromatin Contact Maps Reveals Spatially Active Regions in the Human Genome. Cell Rep 2016, 17:2042–2059.

2. Xiong K, Ma J: Revealing Hi-C subcompartments by imputing inter-chromosomal chromatin interactions. Nat Commun 2019, 10:5069.

